# Changing geographic patterns and risk factors for avian influenza A(H7N9) infection in China

**DOI:** 10.1101/146183

**Authors:** Jean Artois, Xiling Wang, Hui Jiang, Ying Qin, Morgan Pearcy, Shengjie Lai, Yujing Shi, Juanjuan Zhang, Zhibin Peng, Jiandong Zheng, Yangni He, Madhur S Dhingra, Sophie von Dobschuetz, Fusheng Guo, Vincent Martin, Wantanee Kalpravidh, Filip Claes, Timothy Robinson, Simon I. Hay, Xiangming Xiao, Luzhao Feng, Marius Gilbert, Hongjie Yu

## Abstract

The 5^th^ epidemic wave in 2016-2017 of avian influenza A(H7N9) virus in China caused more human cases than any previous waves but the factors that may explain the recent range expansion and surge in incidence remain unknown. We investigated the effect of anthropogenic, poultry and wetland information and of market closures on all epidemic waves (1-5). Poultry predictor variables recently became much more important than before, supporting the assumption of much wider H7N9 transmission in the chicken reservoir, that could be linked to increases in pathogenicity. We show that the future range expansion of H7N9 to northern China may translate into a higher risk of coinciding peaks with those of seasonal influenza, leading to a higher risk of reassortments. Live-poultry market closures are showed to be effective in reducing the local incidence rates of H7N9 human cases, but should be paired with other prevention and control measures to prevent transmission.

## Introduction

A novel avian influenza A (H7N9) virus emerged in spring 2013 in China. The third and fourth epidemic waves of human infections, in the winters of 2014/2015 and 2015/2016 respectively, showed an apparent reduction in incidence compared to spring 2013 and winter 2013/2014 epidemic waves. However, during the winter of 2016/2017, the incidence rose, growing to levels never observed before and reaffirming concerns of a pandemic threat posed by the H7N9 virus (Wang et al. 2017; L. Zhou, Ren, et al. 2017; Uyeki, Katz, and Jernigan 2017). Since 2013, more than 1412 human cases of H7N9 virus have been reported, mostly located in eastern China, with a case fatality risk ranging between 30% and 40% (Yu, Cowling, et al. 2013; Xiang 2016; Z.-Q. Wu et al. 2017). This can be compared to the case fatality risk of H5N1 highly pathogenic avian influenza (HPAI) human infections (53.5%) (Lai et al. 2016). Within China, HPAI H5N1 human cases had a fairly scattered distribution and the majority of cases occurred in the past. In contrast, the annual incidence of H7N9 is higher and distributed in highly populated areas of China (Artois et al. 2016; Bui et al. 2017), making it a virus of particular human health concern.

The H7N9 virus that caused the first epidemic wave in March 2013 originated from multiple reassortment events of avian influenza viruses from domestic poultry and wild birds (Lam et al. 2013) including six internal genes originating from H9N2 strains from chickens. Mainly restricted to Yangtze River Delta in eastern China including urban areas of Shanghai and Jiangsu and Zhejiang provinces in the first wave, the spatial range of H7N9 human cases increased during the second wave along the coast into Guangdong province in southern China (Xiang et al. 2016). Interestingly, recent results from phylogeographic inference suggested that the H7N9 virus sequences of the third wave in central Guangdong likely resulted from local persistence of the virus rather than reintroduction from elsewhere (H. Zhu et al. 2016). The authors suggested that H7N9 has become established and enzootic in different and separate parts of China. The H7N9 virus genotypes of the first wave could have evolved to multiple regional lineages during the second and third waves, reassorting with local avian influenza viruses (Lam et al. 2015; H. Zhu et al. 2016).

While the epidemiological characteristics of human cases are well described, there is poor information on the distribution of H7N9 in its reservoir hosts, chickens. Indeed, humans are not a natural reservoir but an occasional host of H7N9 and the human cases act as sentinels, presumably reflecting the circulation of H7N9 in bird populations (H. Zhu et al. 2016). The surveillance of H7N9 in poultry is difficult because the majority of the virus has a low pathogenicity in chickens (Pantin-Jackwood et al. 2014; Kalthoff et al. 2014) and surveillance can therefore not be based on clinical signs and needs to involve active and targeted sampling. This may change in the future due to the recent evolution of an highly pathogenic strain of H7N9 (W. Zhu et al. 2017; Ke et al. 2017; L. Zhou, Tan, et al. 2017). However, to date, investigations on the spatial distribution of virus reservoirs have remained inconclusive. The distribution of human cases therefore represents the most effective way to study the spatial distribution of H7N9 virus (combined with surveillance findings from birds and environment) and to try gaining knowledge on underlying spatial risk factors associated with the human exposure. Another important aspect of the epidemiology of H7N9 human infections is the role of measures that were taken to close, clean or disinfect live-poultry markets (LPM) to reduce the transmission of the virus along the poultry value-chain, and to reduce human exposure. Many different measures ranging from permanent closures to a weekly day-off with disinfection have been implemented in different provinces and at different times within each epidemic waves. So, market closure, alongside other factors may have an influence on disease risk in space and time. For example, a shift in the spatial distribution of human cases from the urban areas to rural areas may have been related to the implementation of LPM closures in cities after the first wave (Xiang et al. 2016).

Besides market closure and disinfection measures, three sets of factors may have a significant influence on spatial variation in H7N9 incidence.

First, the visits to LPM are clearly the main known risk factor of H7N9 infection at the human case level (Yu et al. 2014; Yuan et al. 2015; J. Wu et al. 2016) and LPM represent a key interface between human, poultry and to some extent, peri-domestic birds. At a higher level, LPM networks may support the persistence of H7N9 virus because chicken movements through the network of LPM and poultry farms may facilitate H7N9 spread and persistence (X. Zhou et al. 2015). In previous studies, we showed that a high density of LPM in some specific areas could regionally increase the risk of H7N9 infection for humans at the market level (Gilbert et al. 2014, 9), which translated into higher risk at the county level quantified in several studies (Fang et al. 2013; Fuller et al. 2014; Li et al. 2015). So, the first set of spatial risk variables, termed “anthropogenic variables”, included the distribution of LPMs and human population density. The latter was included as it may be a good surrogate for some surveillance and reporting bias or for some anthropogenic transmission mechanisms.

Second, 69%-80% of H7N9 human cases of the five epidemic waves reported exposure to live poultry prior to infection, including LPM (52%–60%) and backyard poultry (13%-40%), and these figures remained fairly stable with time (Wang et al. 2017). Whilst the majority of those exposures may correspond to LPM visits, other opportunities for contact with poultry along the production and value chain could take place. For example, poultry workers in Beijing were shown to be at a higher risk of H7N9 infection than the remaining population of the city (Yang et al. 2016). In the general epidemiology of avian influenza emergence, poultry is at the interface between human population and waterfowl, migratory birds and peri-domestic birds (Kapan et al. 2006; Lam et al. 2015; Peiris et al. 2016; Bahl et al. 2016) and may itself become a reservoir if the circulation of avian influenza viruses through the production and value-chain cannot be prevented. Poultry-related variables were found to be significant predictors of H7N9 risk in several previously published studies (Gilbert et al. 2014; Li et al. 2015; Xu et al. 2016; Artois et al. 2016).

However, until recently H7N9 was rarely found in poultry farms through active surveillance and a better understanding of how poultry plays a role in the spread of H7N9 is still needed. During the 5^th^ wave, outbreaks in farms started to be reported with higher numbers and these higher detections may be linked to the emergence of the H7N9 HPAI, which makes passive surveillance more effective through apparent clinical signs. Hence, domestic poultry remains the most likely disease reservoir thus we included a second set of predictor variables, termed “poultry” variables including the density of chickens and ducks, as these may regionally influence the risk of H7N9 virus transmission to humans.

Third, although the most conservative hypothesis remains that human infections would be linked to the circulation of H7N9 in domestic chicken reservoirs with occasional human exposure in LPMs, one can not excluded that wild birds may have taken part in the transmission. The virus precursors of the H7N9 virus in China were found in a wide variety of bird species, wild and domestic (Lam et al. 2013), and avian influenza viruses circulating in wild bird represent a gene pool that may recombine with H7N9 viruses and allow better adaptation and persistence. There is little information on the wild host specificity of H7N9, and data on the distribution of wild bird species is generally fairly coarse, with populations varying strongly according to the season. Because of those uncertainties, proxy variables are required to investigate the possible effect of wild birds on H7N9 transmission to humans, and the third set of predictor variables related to inland water/wetland presence as an indicator of wild water bird distributions.

The aim of this paper was to study the spatial variation of H7N9 incidence in the human population during the 5 epidemic waves in relation to anthropogenic, poultry and wild bird habitat predictor variables on one hand, and in relation to market closure measures on the other hand. The effect of risk factors and market closures had to be analysed separately because market closures were often implemented reactively at the time of epidemics in counties or provinces where incidence was rising. The spatial distribution of market closure measures does in fact correspond well with areas where a high number of cumulative cases were observed over the entire series of epidemics, and constitutes a confounder effect because the distribution of market closure measures would itself be a strong spatial predictor of incidence. Hence, the analysis of market closure measures and other risk factors were carried out separately.

Finally, the analysis was repeated over the five epidemic waves of infection, which allowed the investigation how different predictor variables were linked to H7N9 infection over time, the spatial distribution of repeated re-occurrences, and the year-to-year variation in predictability of H7N9 infections.

## Results

A GLMM models was built to study the association between LPM closure measures and the daily incidence rate (DIR) of H7N9 human cases, by contrasting counties with no measures, counties with measures but before they were taken, and counties with measures and increasing levels of closing days. The GLMM models with the closing status were always more explanatory than intercept-only models based on Akaike information criterion (AIC), and the results of the comparison of DIR according to the closure measures are presented in Figure 1. A detailed analysis of these results shows that, with the exception of waves 1 and 4, the DIR computed for counties with different levels of closing measures were significantly lower than the DIR computed in the same counties before the closure(s) (Before C). The DIR computed for counties which did not implemented any measures but experienced at least one human case in their wave was also comparatively high. The comparison of the weekly numbers of the H7N9 human cases between counties concerned by LPM closures measures and counties free of LPM closures is presented in Fig. 2. Note that in both Fig. 1 and Fig. 2, DIR in counties without any measures was estimated from counties with at least one human case, so the set of countries from which these estimates were derived changed from wave to wave. During all epidemic waves, there remained a fairly high DIR in counties that did not implement any measures, even after the peak of new measures had passed, highlighting that market closure measures only concerned a fraction of counties where they may have been efficient in reducing the number of human cases.

**Figure 1.**
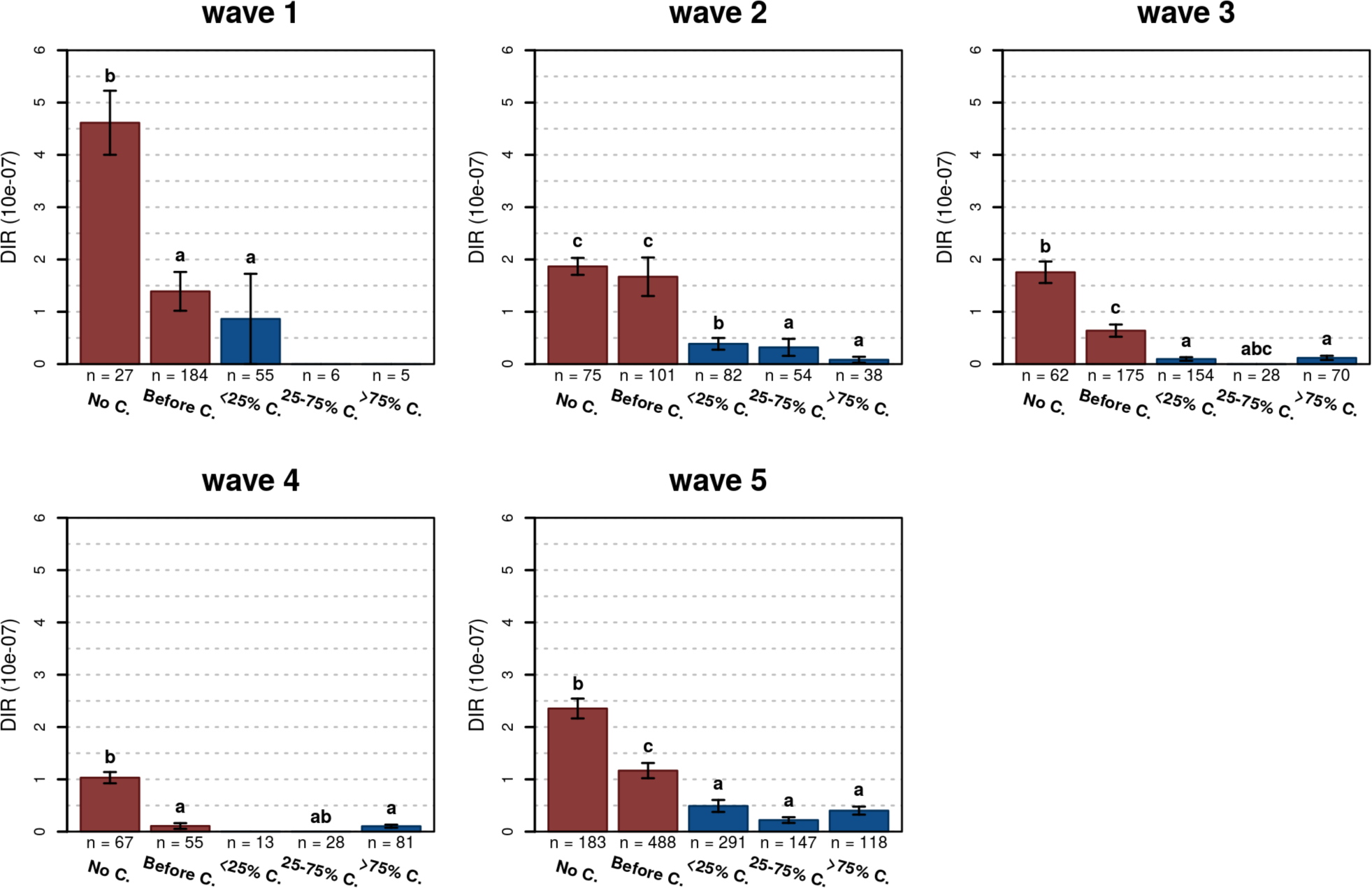
Daily incidence rate estimated during the different epidemic waves in counties with no closing measures but having at least 1 cases during the epidemic wave (No C.), in counties with measures, but before the first closing measure (Before C.) and with different proportion closing days after the first measures.

**Figure 2.**
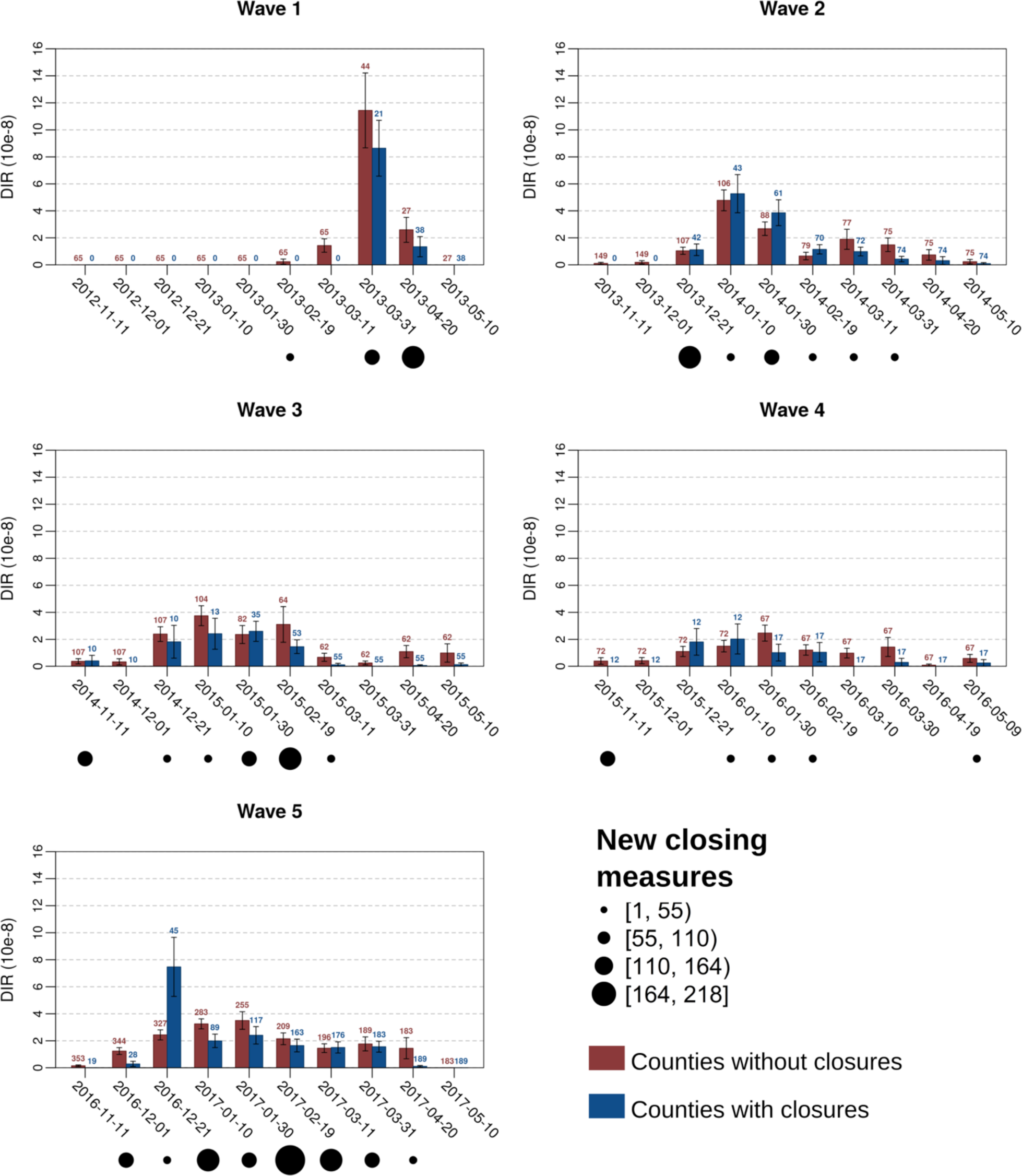
Daily incidence rates in counties with or without closing measures (only including counties with at least one human case over each epidemics). The black dots with varying size are indicative of the number of new closing measures in each 20 days time interval.

A Poisson boosted regression tree (BRT) models was built to predict the DIR of H7N9 human cases as a function of a set of anthropogenic (LPM density, human population density), poultry (poultry density, chicken to duck ratio) and water bird habitat (distance to water, proportion of water in the county) predictor variables. Table 1 presents the relative contribution (RC, a measure of the importance of predictor variables in the BRT models, which quantifies the weighted proportion of use of the variables in the trees) of the different predictor variable of the BRT models in the different epidemic waves. It can first be noted that the RC of anthropogenic predictor variables were generally high (w1 = 40.61%; w2 = 50.12%; w3 = 39.26%; w4 = 17.61%; w5 = 17.94%) but decreased strongly after the third epidemic wave. In parallel, the RC of poultry predictors increased and was greatest in the last epidemic wave (w1 = 10.47%; w2 = 5.83%; w3 = 2.64%; w4 = 28.54%; w5 = 41.83%). In this last epidemic wave, the most important predictor variables were by decreasing order of RC the Chicken to Duck ratio (27.28%), the LPM density (16.04%), the poultry density (14.55%) and the distance to open lakes and reservoirs (6.16%). Fig. 3 presents the BRT profiles of these four predictor variables in the different epidemic waves (the other profiles are provided as supplementary information Fig. 2). The chicken to duck ratio had a significant RC only in waves 4 and 5, when it showed a positive association with incidence up to a ratio of approximately 30. The LPM density profile of wave 5 also showed a positive association with the LPM density, levelling-off at a density of 0.01, and with a relatively similar profile to the other epidemic waves. The 5^th^ wave tended to associate lower incidence with the highest densities (> 0.03), in contrast to previous epidemic waves. The poultry density profile changed gradually over time, with an increasing RC, and the incidence rate in wave 5 is predicted to increase strongly in counties with a very high density of poultry (> 60,000 heads/km^2^). Finally, the profile of the distance to lakes showed a decreasing association, which in the range 0 – 100 km.

**Figure 3.**
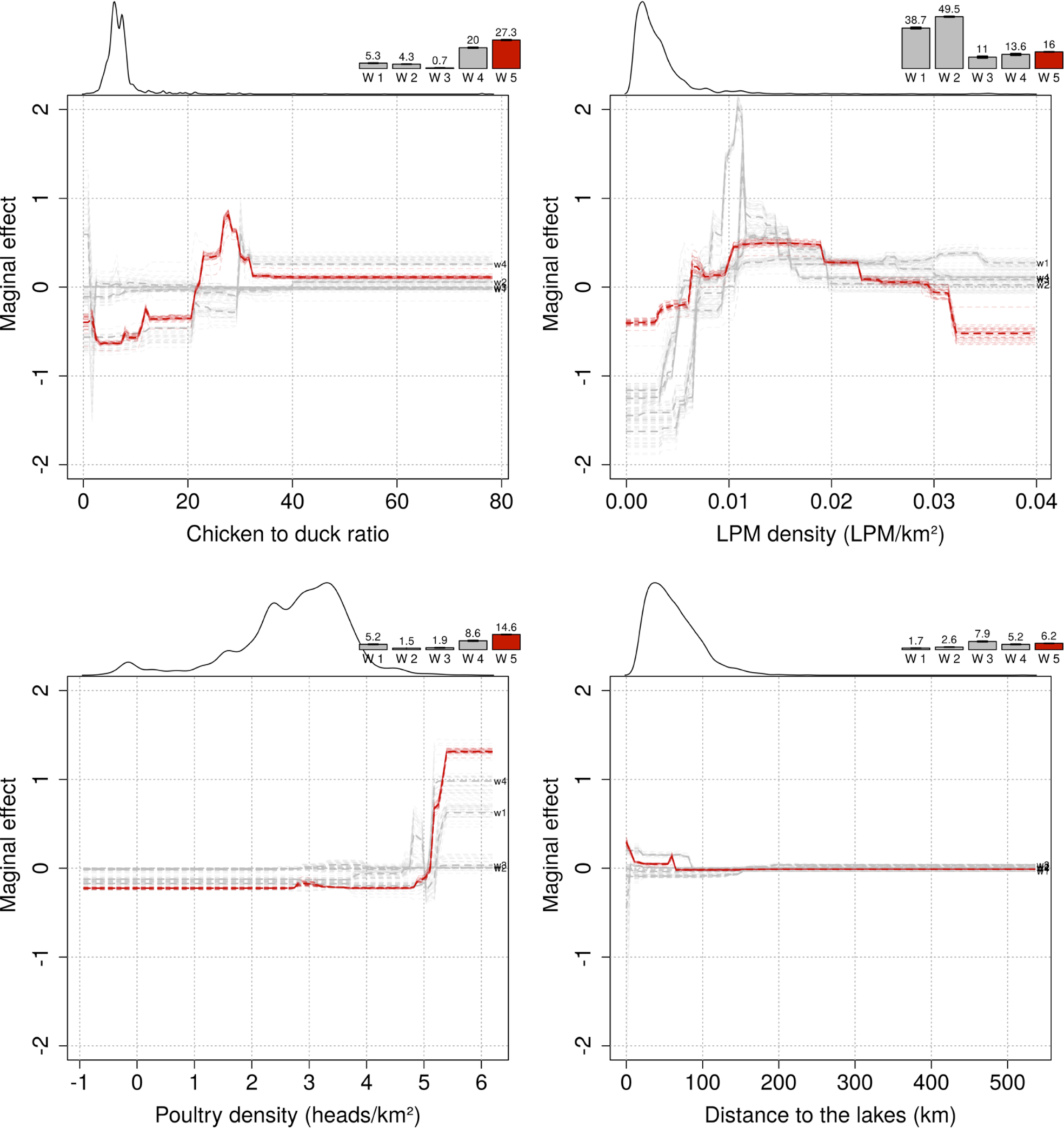
Marginal effect plots of the top-4 predictor variables on the predicted incidence rate, with the change in relative contribution over time indicated by the bars on the top of each plot, showing the increasing relative contribution of the poultry predictor variables. The smoothed line on the top left part of each plot is indicative of the distribution of each variable.

**Table 1.** Relative contribution of the different BRT models across the different epidemic waves

|  | Wave 1 | Wave 2 | Wave 3 | Wave 4 | Wave 5 |
| --- | --- | --- | --- | --- | --- |
| Anthropogenic (sum of RC) | 40.61 | 50.12 | 39.26 | 17.61 | 17.94 |
| LPM density | 38.68 ± 0.76 | 49.48 ± 0.4 | 11.02 ± 0.89 | 13.59 ± 0.77 | 16.04 ± 0.16 |
| Human pop. density | 1.93 ± 0.17 | 0.64 ± 0.03 | 28.24 ± 0.55 | 4.02 ± 0.24 | 1.9 ± 0.06 |
| Poultry (sum of RC) | 10.47 | 5.83 | 2.64 | 28.54 | 41.83 |
| Chicken to duck ratio | 5.29 ± 0.15 | 4.34 ± 0.05 | 0.72 ± 0.11 | 19.96 ± 0.42 | 27.28 ± 0.34 |
| Poultry density | 5.18 ± 0.16 | 1.49 ± 0.04 | 1.92 ± 0.17 | 8.58 ± 0.32 | 14.55 ± 0.11 |
| Water habitat (sum of RC) | 2.21 | 3.94 | 9.75 | 5.95 | 7.67 |
| Prop. of wetlands | 0.55 ± 0.05 | 1.34 ± 0.06 | 1.81 ± 0.12 | 0.8 ± 0.05 | 1.51 ± 0.05 |
| Distance to lakes | 1.66 ± 0.07 | 2.6 ± 0.09 | 7.94 ± 0.31 | 5.15 ± 0.19 | 6.16 ± 0.07 |
| Autoregressive term | 46.7 ± 0.6 | 40.11 ± 0.34 | 48.35 ± 0.77 | 47.9 ± 1.22 | 32.55 ± 0.26 |

The assessment of the BRT models goodness of fit is presented in Table 2, and with the exception of the 4^th^ epidemic waves, the predictability of the models were moderate with cross-validation correlation coefficients within a range from 0.42 to 0.55. In presence/absence term, the models had a good discriminatory capacity with AUC ranging from 0.78 to 0.92 but this decreased over the years (w1 = 0.92; w2 = 0.85; w3 = 0.83; w4 = 0.86; w5 = 0.78). This difference in predictability highlights that it is apparently easier to predict the presence or absence of a human case than it is their number. Epidemic wave 4 was quite specific, longer in time but of lower intensity with a lower total number of human cases than during the other waves, which may explain the lower predictability. The evaluation of the temporal extrapolation capacity of the different models is presented in Table 3. The AUC metrics decrease when a prediction of a given wave is tested for its ability to predict the presence of H7N9 human cases in the following years and AUC values never drop below 0.74.

**Table 2.** Goodness of fit metrics of the BRT models across the different epidemic waves

|  | Pearson Corr. Coefficient |  |  | AUC |  |
| --- | --- | --- | --- | --- | --- |
|  | Training | Training_Auto | CV | Training | Training_Auto |
| wave 1 | 0.785 ± 0.012 | 0.552 ± 0.002 | 0.477 ± 0.015 | 0.923 ± 0.001 | 0.907 ± 0.002 |
| wave 2 | 0.753 ± 0.004 | 0.326 ± 0.007 | 0.548 ± 0.013 | 0.85 ± 0.001 | 0.849 ± 0 |
| wave 3 | 0.582 ± 0.011 | 0.495 ± 0.004 | 0.42 ± 0.013 | 0.831 ± 0.003 | 0.814 ± 0.001 |
| wave 4 | 0.44 ± 0.012 | 0.293 ± 0.006 | 0.259 ± 0.008 | 0.856 ± 0.001 | 0.832 ± 0.001 |
| wave 5 | 0.61 ± 0.002 | 0.527 ± 0.002 | 0.447 ± 0.009 | 0.778 ± 0.001 | 0.755 ± 0 |

**Table 3.** Cross-predictability of the models trained with the different epidemic waves applied to the others, as measured by the AUC.

|  | Applied to |  |  |  |  |
| --- | --- | --- | --- | --- | --- |
|  | wave 1 | wave 2 | wave 3 | wave 4 | wave 5 |
| Predictions of |  |  |  |  |  |
| wave 1 | 0.91 | 0.81 | 0.78 | 0.84 | 0.79 |
| wave 2 | - | 0.85 | 0.78 | 0.83 | 0.76 |
| wave 3 | - | - | 0.82 | 0.82 | 0.74 |
| wave 4 | - | - | - | 0.83 | 0.75 |
| wave 5 | - | - | - | - | 0.76 |

Figure 4 shows the distribution of the top-three predictor variables (Live-poultry market density, poultry density and chicken to duck ratio) in relation to the distribution of the past and last epidemic wave. The RGB composite plot (Fig. 4a) highlights areas where all three predictor variables were high and where H7N9 persisted over time (Fig. 4b), as indicated by the number of years during which each county experienced at least one human case. First, in a large areas surrounding the Yangtze River delta, poultry production is largely dominated by chicken production to the north of the Taihu Lake, high live-poultry market density to the east of the Taihu Lake on the urban areas of Wuxi, Suzhou and Shanghai and includes several small hotspots of high poultry production in a large area that surrounds the Taihu Lake. Second, the RGB composite plots highlight three additional urban areas with high LBM density and high poultry density: the Guangdong province, the Tianjin and the Beijing urban areas and the Chongqing urban area. These different areas highlighted in the RGB maps visually correspond to areas of high H7N9 re-occurrence displayed in Fig. 4b. Indeed, the count of number of years with at least one human case helps to visualise the distinction between counties with repeated reoccurrences from counties with sporadic infections. These areas include southern Jiangsu, Shanghai and northern Zhejiang provinces, as well as Guangdong counties located around Hong Kong but to a lesser degree than the areas in and around Shanghai. Fig. 4c highlights that the spatial pattern of wave 5 showed a marked geographic expansion from these previous hotspots of persistence, with a 90 counties reporting H7N9 for the first time (50.85% of the total number of counties infected in wave 5). One can also measure why live-poultry density was a lower predictor in wave 5 than in previous waves, as these newly infected counties do not match green areas depicted in Fig. 4a.

**Figure 4.**
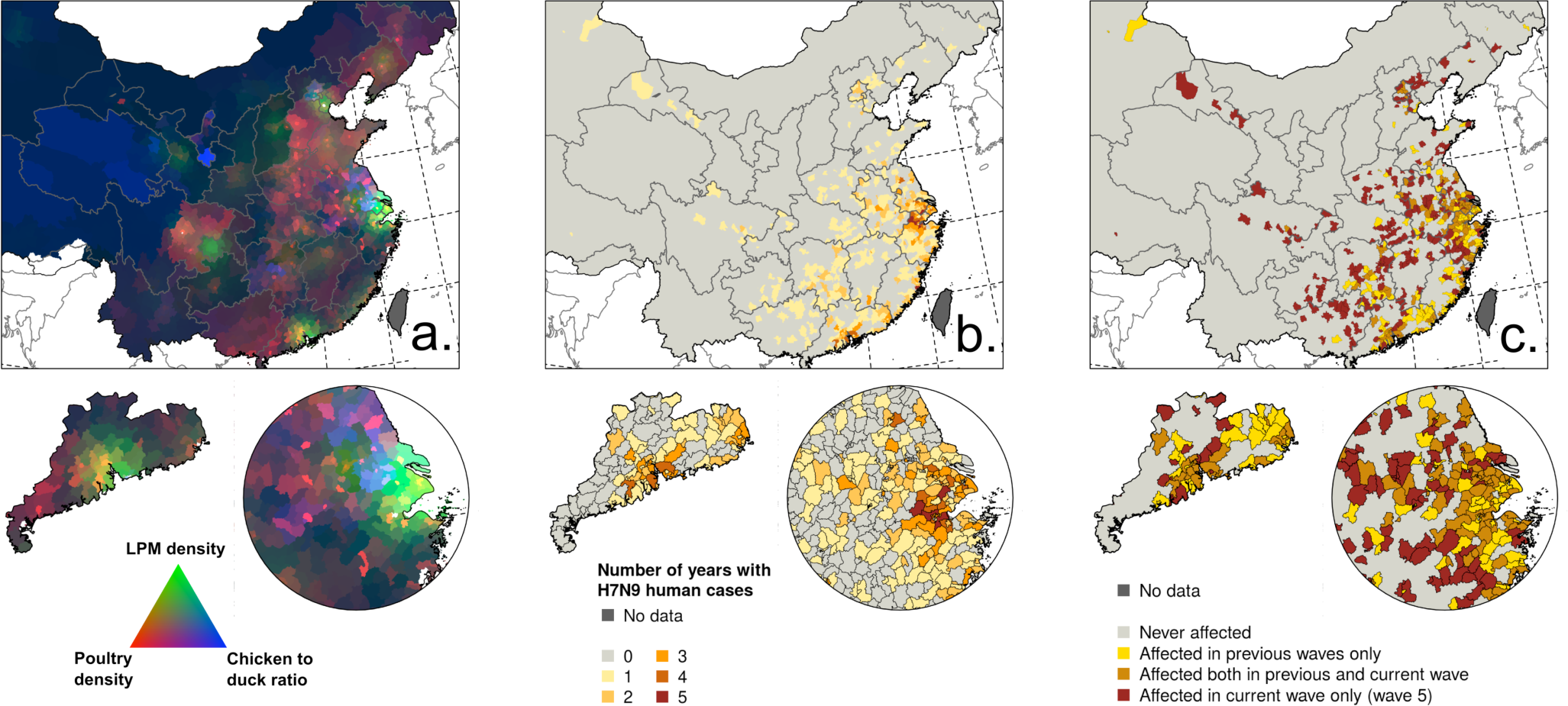
Distribution of predictor variables and H7N9 infections. A. Red-Green-Blue visualisation of poultry density (red), live-poultry market density (green) and chicken / duck ratio (blue), with dark areas corresponding to low values in all three predictors, and white areas to high values in all three predictors. B. Number of years with at least 1 human case per county. C Distribution of the 5th wave of human infections over previous ones.

The heat maps presented in Fig. 5 show that until now, the majority of H7N9 human cases have taken place in February to March (Fig. 5B) with a latitudinal gradient. The seasonality of common influenza A infection shows different levels of seasonality in China (Fig. 5C), with the province north of 34.1 degrees showing a much stronger annual winter seasonality of infection than more southern provinces, with a peak in December – February. By comparing figure 5B to 5C, once can visualise that so far, the peaks of H7N9 and seasonal influenza have not yet been strongly coinciding in space and time. However, a geographic range expansion of H7N9 infections in the northern provinces, keeping its current seasonality, would bring the H7N9 and seasonal influenza incidence peaks to coincide much more extensively.

**Figure 5.**
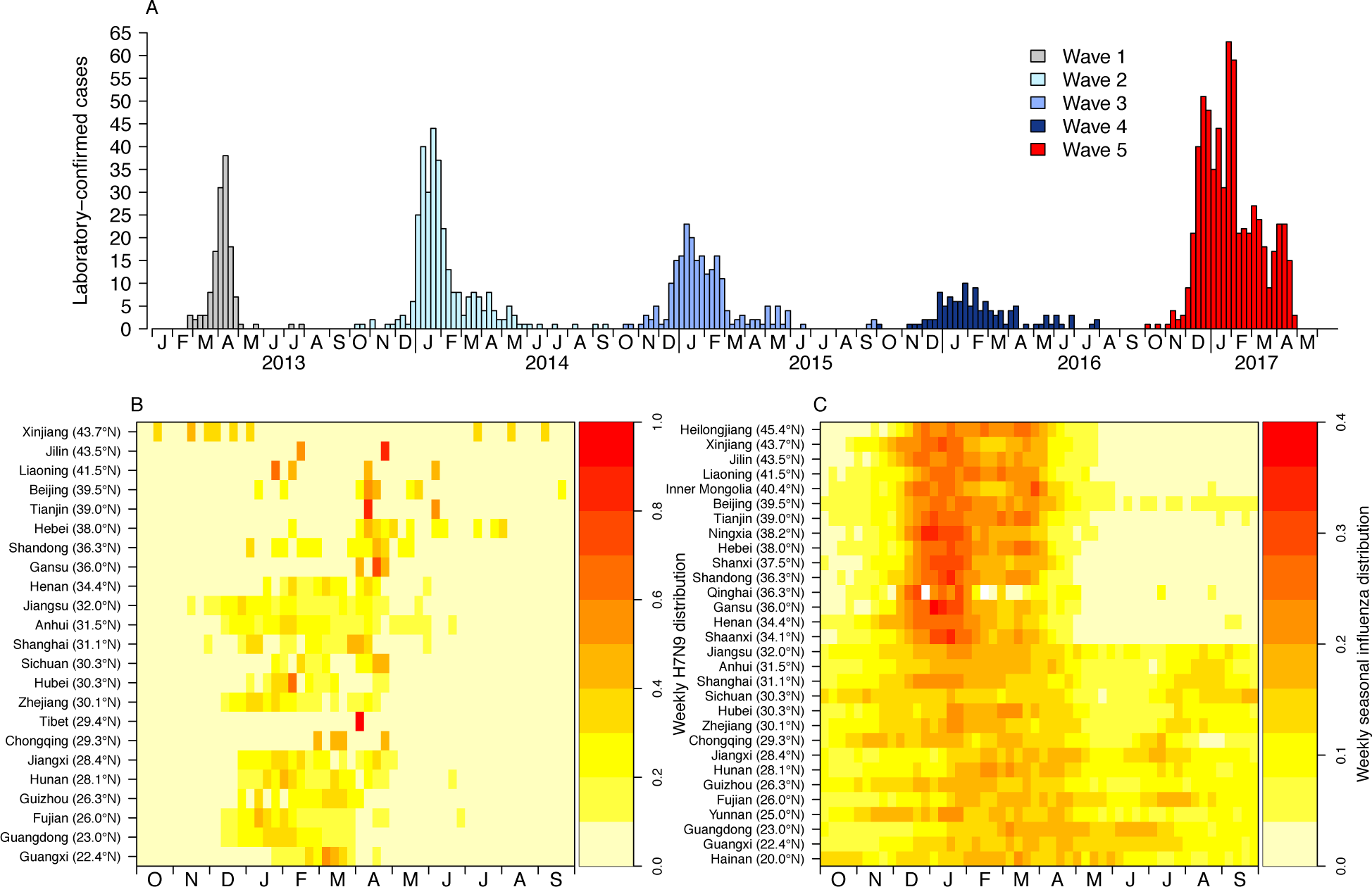
Seasonality of H7N7 infections in comparison to seasonal influenza. A. epidemic curve for H7N9; B seasonality for H7N9; C seasonality for seasonal influenza.

## Discussion

The results of our spatial models demonstrate a significant shift over time from anthropogenic to poultry predictor variables linked to H7N9 human cases. This shift was already apparent in the 4^th^ epidemic wave, although fewer human cases were reported. More specifically, the predictive power of poultry variables increased over time and was greatest in the last epidemic, pointing to areas with very high chicken densities and high chicken to duck ratios. A recent study on H7N9 human cases showed an increase in semi-urban and rural cases in the last wave, and a comparatively higher number of middle-aged cases (Wang et al. 2017). However, apart from the overall increase in cases, the study did not suggest any other major epidemiological differences, and other authors made similar observations when comparing waves 1-4 (Wang et al. 2017; Xiang et al. 2016; Xiang 2016). Our results do not contradict the observation of a higher number of human cases in peri-urban and rural areas, because high poultry production regions are typically located in peri-urban and rural settings. But they strongly support the hypothesis that the H7N9 virus may have spread in the chicken reservoir much more extensively in the last two epidemic waves than previously, with a particularly marked geographical range expansion in the last epidemic wave. This observation based on human case can be linked to the emergence of HPAI H7N9 that was reported early 2017 in southern China (W. Zhu et al. 2017). Recently published results showed that human cases of HPAI H7N9 were already found beyond Guangdong, in Hunan and Guangxi in early 2017 (L. Zhou, Tan, et al. 2017). In parallel, there was a comparatively higher number of reports of H7N9 positives found in poultry farms this year in comparison to previous epidemic waves, including reports of HPAI H7N9 in northern China, in Tianjin (FAO Empres 2017). The precise role of the gain in pathogenicity on the range expansion of H7N9 is yet unclear, as of the main mechanisms of transmission along the poultry production and value chain networks. However, the fact that such a range expansion took place in parallel to the emergence of a highly pathogenic variant can hardly be coincidental.

It should be borne in mind that the measure of predictor weights in the model is relative, i.e. the sum of relative contribution equals to 1, so if poultry variables become better predictors of H7N9 incidence in human, the RC of other variable would decrease, even if their effect on the predicted incidence remained fairly constant. This seems to be the case for the LPM variable, as the BRT profiles remained fairly stable, suggesting that the role of LPMs in the transmission may have remained important, and adding up with the increasing contribution of the poultry predictors to lead to the highest incidence observed in the 5^th^ wave. In other words, the contribution of LPMs may have remained high, but its combination with increasing transmission along the poultry production and value chains may be responsible for the geographical range expansion and higher incidence of the 5^th^ wave.

Although some of the highest incidences were observed along Taihu Lake, the predictive capacity of water bird-related predictor variables appeared to have a much lower influence on the predicted incidence than anthropogenic and poultry variables. Wetlands constitute favourable ecosystems for the emergence of new avian influenza viruses, especially when intensive poultry farming is taking place in nearby landscapes, forming ideal interfaces for wild-domestic and domestic-wild transmission. Many interfaces combining wetlands, intensive poultry farming and rice paddy fields are present in south-eastern China and may have played a role in the initial emergence of the H7N9 virus in the Shanghai area (Liu et al. 2013). However, as the virus spread more abundantly in the domestic chicken reservoir in the following epidemic wave, the contribution of wild birds to overall disease circulation may be fairly low, which is reflected by the low relative contribution of the water bird proxy variables.

The predictive capacity of the incidence models was only moderate, and this can naturally be explained by the fact that these spatial models didn’t account for the variability in incidence linked to market closure measures. This is confirmed by the fact that the predictions of presence/absence were generally better, because presence cannot be influenced by market closure measures (as they followed human cases rather than preceding them), and few counties implemented market closure measures in the absence of human cases.

The geographical range expansion and increases in incidence of human cases in the 5^th^ wave of H7N9 brings serious human health concerns. First, repeated human infection by avian influenza viruses increase the chances of adaptation to improved human to human transmission. Second, the provinces affected by earlier H7N9 epidemic waves do not have a strong seasonal influenza A peak in January and February (Yu, Alonso, et al. 2013) that matches the peak of H7N9 cases (Fig. 5). However, if the H7N9 virus continues to expand its range northward, in areas with a strong influenza A peak in January and February, there will be a higher chance of local coincidence of peaks of incidence between human cases of H7N9 and seasonal influenza A virus. This may enhance the chances of co-infections that could lead to the emergence of reassortants with the capacity to easily transmit between humans. Third, the extent of the geographical range of the expansion is not yet fully known and in the absence of new measures, it may spread further within China, and internationally through poultry value-chains.

Preventing human infections has so far mostly relied on market closures, and our results based on the 5 waves tend confirm the efficiency of the reduction reported in previous studies. For example, (2014) showed that the closure of LPM reduced the mean daily number of infections by a factor ranging between 97% and 99% during the first epidemic wave in 4 cities of central-eastern part of China (Shanghai, Hangzhou, Huzhou, and Nanjing). However, as highlighted in Fig. 2 showing the perpetuation of epidemics despite market closures, with relatively high incidences in counties with no measures, the implementation of LPM closing measures and their efficiency is fairly heterogeneous in space and time and, with the exception of permanent closure, was most often implemented reactively in areas directly affected by H7N9 human cases. Live-poultry market closing measures may have a direct effect on the local circulation of the virus in the market networks and value-chain and on the human exposure. However, the poultry trade network in China is extensive and complex and poultry sold in a market may come from many different sources distributed throughout the country (Martin et al. 2011; Soares Magalhães et al. 2012; X. Zhou et al. 2015). For example, an interview of 25 live poultry traders in southern China Magalhaes et al. (2012) showed that in February, 4 LPM were linked to 122 poultry sources with an average distances between the poultry source and LPM of 803 km. Live poultry traders could recoup the loss of sales points by enlisting nearby or more distance areas that were less concerned by LPM closing measures. An interesting aspect of the study of Magalhaes et al. (2012) is that the geographic extent of the poultry source/LPM network appears highly dependent of the period of the year and maximal in February and around the Chinese New Year festivities. More recently, a similar analysis was done in Yangtze River delta and surrounding areas (central-eastern China) and showed selected LPM from Shanghai, Nanjing, Hangzhou, Huzhou, Hefei and Chuzhou had large catchment areas with poultry being brought from adjacent provinces (from January to April) (X. Zhou et al. 2015). So, unless LPM is paired with poultry movement controls or pre-movement testing policies, the measure may be effective locally but may also have implications on the risk of disease spread to other areas. A second aspect is that the reactive nature of market closure in the most severely affected areas necessarily implies a significant delay. By the time human cases become detected in sufficient numbers to trigger a reactive market closure decision, the disease has had many opportunities to spread through the market networks to other areas not necessarily concerned by the measure. This probably explains why many human cases continue to be reported in counties with no measures for several weeks after the peak of market closure measures.

## Conclusion

A major shift was observed between the predictor variables of H7N9 human cases during the course of the five epidemic waves, with poultry predictor variables becoming more important than anthropogenic predictor variables, particularly in waves 4 and 5. This result strongly supports a recent and significant geographical expansion of H7N9 viruses, and their circulation in the poultry reservoir. This could have been paired with a higher pathogenicity and result in the observed increase in frequency of reports in poultry farms.

The current range expansion of H7N9 to more northerly latitudes may increase the chances of H7N9 peaks coinciding in both space and time with those of seasonal influenza A infection, leading to higher risk of co-infections and reassortments.

LPM closure measures appear to be effective in reducing the daily incidence rates of H7N9 human cases in counties where they are implemented, especially when these measures are permanent, or implemented for a sufficiently long period. They do not, however prevent the reporting of human cases in other areas, so their impact appears mostly to be local. In addition, such measures drive poultry trade elsewhere, e.g. in peri-urban areas or the countryside, with related human exposure risk.

Reducing H7N9 circulation in poultry and humans may require significantly more preventive and control measures than those currently in place. The following, for example, could be considered: i) pro-active rather than reactive market closure measures in areas and periods predicted to be at high risk, based on restrospective space-time modelling of previous waves of infections, ii) better surveillance, prevention and control of H7N9 along the poultry production and value-chain networks, iii) poultry movement bans or pre-movement testing policies, and iv) the development and use of vaccination to prevent or reduce H7N9 circulation in poultry, following the experience of China’s mass-vaccination of poultry against the HPAI H5N1.

## Material and methods

### Data

#### H7N9 human cases and seasonal Influenza

All confirmed H7N9 human cases during the period from March 2013 to 18^th^ April 2017 were analysed. These were surveillance data from the national system centralised by the China Centre for Disease Control and Prevention (China CDC). A detailed description of case definitions, surveillance for identification of cases, and laboratory testing for A H7N9 virus have been provided elsewhere (Cowling et al. 2013; Yu, Cowling, et al. 2013; Qin et al. 2015). For each case, the information about place of residence and date of onset of symptoms were used. Indeed, this study predicted that exposure lead to symptoms within 6.5 days (95% confidence interval: 5.9, 7.1), so 6.5 days have been subtracted from the date of onset of symptoms to estimate the dates of first contact with the virus (Virlogeux et al. 2015). Finally, in order to compare the seasonality of H7N9 human cases with that of common influenza A in space and time, we used data from the national sentinel hospital-based influenza surveillance network, providing the weekly proportion of laboratory-confirmed influenza cases by virus type from specimens tested in 30 Chinese provinces during the period January 2013 – March 2017. More information on the sentinel network from which these data were derived can be found in Yu et al. (2013).

#### Live poultry markets and closing measures

A database recording the location of LPMs, type of market closure measures, with starting and end date, implemented since the first wave was compiled by the authors. The database was initially assembled by combining i) data from the official website of the Ministry of Agriculture of China and agricultural bureaus at province and prefecture level, ii) a database of points of interest from the official gazetteer issued by the National Administration of Surveying, Mapping and Geoinformation, and iii) several unpublished data sources obtained through data mining, internet searches and direct contacts with provincial Agricultural bureaus.

A database recording the type, starting date, end date and location of market closure measures that were implemented since the first wave was compiled by the authors. A total of 38 types of measures over different periods of time were implemented in response to the H7N9 human infections, as listed in SI Table 1, and the market closure were implemented at the county or district level. The database was used in two different ways. First, data on permanent market closures was used to update a yearly distribution of LPM locations, by annually removing the permanently closed markets out of the 8,943 retail and wholesale poultry market locations of the full LPM database. Second, the range of closure measures was reclassified according to the proportion of closing days (from 1 day per month to permanent closure), and used to estimate different types of incidence, as detailed in the analysis section below.

#### Spatial predictors

The first set of predictor variables included the LPM density (LPM/km^2^) and human population density (people / km^2^). However, some counties do not have LPMs but the people living there may easily go to surrounding counties. In addition, the role of LPM density may act at a higher level by providing a network of markets through which the disease could spread and persist. So, the LPM density was computed by means of a Gaussian smoothing kernel function with the optimal bandwidth found by Gilbert et al. (2014). For each wave, the LPM density estimate only included markets from counties without permanent LPM closure, hence resulting in a different LPM density distribution per epidemic wave. For human population, we used the human population density from the 2010 census (China Data Center).

The second set of predictor variables included poultry, chicken and domestic duck density from a recently published database with the reference year 2010 (Artois et al. 2016). This new data set was produced using the Gridded Livestock of the World methodology (Robinson et al. 2014; Nicolas et al. 2016) applied to an extensively improved data set compiled by the authors. However, at the county level, there was a very high correlation between duck and chicken density. So in order to reduce collinearity and make the results more easily interpretable, we built two alternative predictor variables: the poultry density (chicken + duck heads / km^2^) and the chicken to duck ratio (chicken heads / duck heads), which were found to be much more independent.

The last set of predictors dealing with water bird habitat included two variables. First, the distance to the largest lakes and reservoirs (km), which represents the distance between the county centroids and the nearest lakes (area ≥ 50 km^2^) or reservoirs (storage capacity ≥ 0.5 km^3^) (Lehner & Döll 2007), and second, the proportion (%) of county covered by wetlands, which was derived from the hybrid wetland map for China (Ma et al. 2012).

Climatic data were not tested in this analysis, for a number of reasons. First, the mechanism by which they may influence human cases is unclear. For example, influenza A human infections generally peak in January-February in Northern China, and in April to June in the southernmost regions (Yu, Alonso, et al. 2013), which does not fit the peak of H7N9 cases in southern China. Seasonality of H7N9 poultry infections is unknown, but apparent for avian influenza poultry infections by HPAI H5N1 viruses. However, continental-scale climatic spatial variables failed to provide robust HPAI H5N1 models in comparison to host-related variables (Dhingra et al. 2016). Second, spatial model pools human cases data over a relatively long period of times with changing conditions that would be more fully taken into account in spatio-temporal models, alongside other variables varying in time such as poultry trade, for example, but these investigations go beyond the scope of the present study.

### Analyses

The five H7N9 epidemic waves had different durations and starting dates. So, in order to make the estimates of incidence comparable, the use of a similar epidemic start and end date for all waves was not justified. Using an epidemic period determined by the first and last case would also be somewhat misleading because the difference between the minimum and maximum is a very sensitive indicator of a distribution spread. So, the duration of each epidemic was set as the period separating the 5^th^ from the 95^th^ percentiles of the days of onset of illness in each wave.

Our first set of analyses focused on the impact of LPM closure on incidence. Daily incidence rates (DIRs) were computed and compared at county level before and after different sets of measures were put in place for the 5 epidemic waves of H7N9. The DIR was defined as the number of new human cases per county population per day over the timespans (number of cases / (population * number of days)). These DIRs were estimated according to different levels of market closures. First, we estimated the DIR in counties without market closures over the full duration of the epidemic wave, but where at least one human case was notified during the epidemic wave (NoC). Second, the DIR was estimated in counties where there was at least one closing measures during the epidemic wave, but in the period preceding the implementation of the first one (BeforeC). Third, DIRs were estimated in counties after the implementation of the first measure and until the end of the epidemics, and we contrasted different levels of closures in that period: low (< 25 % of closing days), intermediate (25 – 75 % of closing days) and high (> 75% of closing days). A generalized linear mixed effect models (GLMM) followed by a multiple comparison procedures (Bretz et al. 2011) was used to compare the DIRs according to the different types of closure. The GLMM models were formulated with Poisson distribution taking into account the county level as random effects (two observations of the same county, before and after the closure, may be considered separately in the models and represent a bias to the assumption of independence of observations). All DIR estimates were done only in counties where the H7N9 virus was reported at least one time over the 5 epidemic waves, such as to exclude the counties in regions that were never reported as infected. Furthermore, DIR estimated from very short period of times may have very variation due to the stochasticity of case reports. So, the GLMM model only included DIR estimates out of durations higher than 20% of the full epidemic wave duration.

Our second set of analyses involved the development of Poisson Boosted regression tree (BRT) models to predict the daily incidence rate of H7N9 virus in human population as a function of the set of predictor variables. The models were developed using the number of human cases as dependent variable with an offset term corresponding to the product of human population by the duration of the epidemic. We build one model per epidemic wave to be able to compare the effect of predictor variables, and to assess the predictive capacity from one wave to another. Each epidemic wave model was developed using a 4-fold cross-validation procedure (Elith, Leathwick, and Hastie 2008) and the Pearson correlation coefficient between the predicted and the observed response was used as a measurement of the predictive capacity of models. The contribution of each predictor variable to the model was quantified in two ways. First, the relative contribution (RC) of each variable is a measure of its overall importance in the model and corresponds to the number of times a variable is selected for splitting the regression trees, weighted by a factor on the quality of the splitting (Friedman and Meulman 2003). This number is expressed in relative frequency so the sum of RC of the individual variables should equal to 1. Second, for each variable and model, we estimated the partial dependence plots, or BRT profiles, which provide a graphical description of the predicted effect of a predictor variable on the response (the incidence rate) after accounting for the average effects of all other predictor variables in the model (Elith, Leathwick, and Hastie 2008).

The presence of spatial autocorrelation in the model residuals was tested using spline correlograms (BjØrnstad and Falck 2001) and the approach of Crase, Liedloff, and Wintle (2012) was used when autocorrelation was present in the model residuals. An autoregressive term containing a spatial average of the initial model residuals is build, and added as predictor variable of a new model. The goodness of fit metrics were both estimated from the models without the autoregressive term and with the autoregressive term set to zero. Finally, in order to account for sources of uncertainty in data splitting for the cross-validation, the analysis was bootstrapped with 30 independent BRT run for a total of 120 cross-validations (30 runs × 4-folds) per wave. The six variables, namely the LPM density, the human population density, the poultry density, the chicken to duck ratio, the distance to open lakes and reservoirs, and the proportion of land covered by water were tested in all models.

In order to test the capacity of the models to discriminate between the presence and the absence of human cases at county scale, the predicted incidence rate was also converted into a probability of having at least one human case in the county. As the population per county (*n*) was high and the mean probability of having a H7N9 human cases (*p*) is very low: B(n, p) ~ P(n*p). Therefore, the probability of having at least one human case per county was estimated with a Binomial distribution as following:

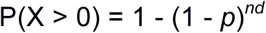

where *nd* is the population times the number of days in the epidemic duration; and *p* is the incidence rate predicted by the Poisson BRT model.

Finally, we also wanted to evaluate the temporal extrapolation capacity of the BRT models, and each model trained with the data of a given epidemic wave was evaluated in its ability to predict the H7N9 human cases from the following epidemic waves. The observed presence or absence of human cases at the county level in a given epidemic wave was compared to the predicted probabilities of human case present of the previous years’ model with the area under curve of the ROC plot.

## Acknowledgements

We thank Shuanbao Yu from Chinese Center for Disease Control and Prevention and Wanqi Yang from Wuhan University for assistance in data collection. The findings and conclusions in this report are those of the authors and do not necessarily represent the views of the Food and Agriculture Organization of the United Nations.

